# Mouse lemurs’ and degraded habitat

**DOI:** 10.1101/216382

**Authors:** Simon Knoop, Lounès Chikhi, Jordi Salmona

**Author notes:** Corresponding authors: Simon Knoop:, Jordi Salmona.

## Abstract

Madagascar is known for its unique biodiversity including its endemic primates, the lemurs. This biodiversity is threatened by deforestation, forest degradation and anthropogenic disturbances. Several mouse lemurs (genus *Microcebus*) have been shown to cope with habitat disturbances and degradation. However, there are 24 recognized mouse lemur species living in very different habitats, and it is not clear whether all these species respond similarly to forest degradation. Here, we review the literature on mouse lemur use of degraded habitat. We further question whether mouse lemurs show variation in degraded habitat use, with respect to forest type, conservation status and distribution range. We show that data on degraded forest (DF) use is available for 14 species and geographically aggregated in a few locations. However, data are scarce for most species, and lacking for almost half of the currently recognized species. Our results however confirm that most mouse lemur species are able to cope with, but do not necessarily respond positively to habitat degradation. We found no variation in degraded habitat use, with respect to forest type, conservation status and distribution range. However, we identified food resources availability, understory structure, predation, and tree hole availability to be the most frequently invoked factors potentially influencing DF use. The relative frequency of these four factors vary among forest types suggesting that differences may exist but still require research efforts for ecological and environmental differences among regions to be fully understood.

**RESEARCH HIGHLIGHTS:**

- Little differences in the use of degraded forest (DF) between forest types, distribution ranges or conservation status.
- Varying factors potentially affecting DF use, such as food resources, forest structure, tree hole availability and predation.

## INTRODUCTION

Madagascar is considered one of the world’s “hottest” biodiversity hotspots due to its exceptional biodiversity and the high level of threats this diversity faces (Goodman & Benstead, 2005; Myers, Mittermeier, Mittermeier, Da Fonseca, & Kent, 2000). Home to *ca.* 110 currently recognized lemur taxa (Louis Jr. & Lei, 2016; Mittermeier et al., 2014; Setash, Zohdy, Gerber, & Karanewsky, 2017), Madagascar harbors the second-highest primate diversity of all countries and the highest proportion of primate endemism (Mittermeier et al., 2010). Mouse lemur habitats and population sizes are decreasing, while their level of threat is rising, mainly from deforestation, forest degradation, and poaching (IUCN, 2017; Schwitzer et al., 2013; Schwitzer, Mittermeier, et al., 2014). Since 2014, 18 out of 24 recognized mouse lemur species are considered threatened (IUCN, 2017; Schwitzer et al., 2013). This high proportion of threatened species is not surprising if we consider the high rate (>50% between the 1950’s and 2000) of forest loss in Madagascar (Schwitzer, Chikhi, et al., 2014). However, they contrast with data suggesting that some species of mouse lemurs are able to use degraded habitat (Ganzhorn, 1995; Mittermeier et al., 2010). Mouse lemurs are commonly observed in degraded forest (we use here a large definition including partially logged, partially deforested, partially cultivated, regenerating forest, but not completely denuded landscape, cf. Methods section for details) (Herrera, Wright, Lauterbur, Ratovonjanahary, & Taylor, 2011; Miller et al., submitted; Randrianambinina, Rasoloharijaona, Rakotondravony, Zimmermann, & Radespiel, 2010), rural areas (Deppe, Randriamiarisoa, Schütte, & Wright, 2007; Ganzhorn, 1987), and in garden environments (Irwin et al., 2010). Aside these evidences, mouse lemurs are forest-dwelling species, and depend on forest for survival (Ganzhorn & Schmid, 1998; Karanewsky & Wright, 2015). Thus, DF might only harbor sink populations. Understanding the use of DF by mouse lemurs may therefore be crucial to their conservation (Schwitzer et al., 2013).

Dry and humid forest species typically have a different diet (Kappeler & Rasoloarison, 2003; Radespiel, 2007), and dry forests generally harbor higher population densities than humid forests (Randrianambinina et al., 2010; Setash et al., 2017). In addition, western dry and eastern humid regions harbor contrasting climatic conditions and climatic extremes that may have led to the development of independent unique resource use strategies (Génin, 2008, 2010; Kobbe & Dausmann, 2009). We therefore ask the following question (Q1): *“Do mouse lemurs vary in their responses to DF in humid and dry forests?”*

Mouse lemur species show a large diversity of distribution range size. Species with large distribution (e.g. *M. murinus*) show high seasonal variability in feeding behavior and high colonization ability (Radespiel, 2016). Contrastingly, other species are stuck in small areas for yet not always clear reasons. We therefore ask the following question (Q2) *“Do mouse lemur species with different distribution ranges vary in their responses to DF?”*

Finally, conservation status is primarily based on population and distribution trends as well as on threats faced by the species (IUCN, 2012). In other words, it summarizes a large panel of factors that may be involved in the ability of mouse lemur species to use DF. We therefore question (Q3) if *“species with different conservation status vary in their responses to DF”*.

This paper reviews mouse lemur DF use and investigate the three abovementioned questions (Q1-3). Finally the present work emphasizes the most commonly invoked and reported factors potentially affecting DF use.

## METHODS

We searched “JSTOR”, “Science Direct”, “Wiley”, “Springer Link”, and “Google Scholar” databases as well as all issues of “Lemur News*”* and *“*Primate Conservation*”* for “*Microcebus*”, “adaptation”, “habitat use” and “habitat degradation”. From identified papers we subsequently searched for species, sex, forest type (dry, humid) and degradation level (cf. classification below), type of degraded habitat use reported: positive, neutral and negative responses to DF and factors invoked (diet, habitat characteristics, sleeping sites, seasonal variation in habitat use, daily torpor/hibernation, territoriality, home range size, competition/coexistence). All studies reporting the presence or absence of mouse lemurs in DF and/or assessing mouse lemur habitat or diet preferences were considered. Review papers reporting information from case studies were not considered.

To compare degradation levels and mouse lemur responses to DF described in different manners in the considered studies, we categorized them, based on the terminology used by the authors. The following terms were considered for primary forest: primary, pristine or natural forest, unexploited forest, undisturbed forest, intact forest, continuous canopy, high density of large trees, high tree species diversity, and absence of human activities. For DF, we considered secondary forest, lightly, moderately, severely degraded or disturbed habitat, forest edges, *savoka* (*i.e.* transitional secondary vegetation after abandonment of agriculture (Radespiel et al., 2012)), forest harboring human activities such as logging, mining, charcoal production, cattle grazing, and fire or traces of fire. For cultivated areas, we considered plantations or areas of slash-and-burn agriculture *i.e. tavy*. Open sites, grassland or savanna were categorized as grassland. A factor putatively influencing DF use was considered when specifically investigated in a particular study. Studies conducted by the same researchers, on the same species, in the same study area and presenting similar results were pooled in a “study cluster” (cf. Table 2). Each “study cluster” was treated as one study in the evaluation. Results on more than one species reported in a single study were considered independently. From now on, all single studies and study clusters are called “studies” without distinction.

The taxonomy of mouse lemurs was subject to regular changes within the last decades (Hotaling et al., 2016; Lei et al., 2015; Rasoloarison, Weisrock, Yoder, Rakotondravony, & Kappeler, 2013; Schwitzer et al., 2013; Thiele, Razafimahatratra, & Hapke, 2013). Hence, former species names were modified to fit the latest taxonomy. Both current and original species names are mentioned in Table 2.

We retrieved the size of each species’ distribution (extent of occurrence, EOO) from the IUCN red list database (IUCN, 2017). Since there is a large uncertainty in the way EOOs are drawn, we assigned each EOO to one of three categories to distinguish small and large distribution range species: “small” (for microendemic species with very few localities or an area of less than 2100 km²), “large” (for species with large distributions of more than 8350 km², i.e *M. murinus*, *M. griseorufus* and *M. myoxinus*), and “medium”. This category comprises the remaining species that do not fall in any of the other two categories (with distributions between 2100 and 8350 km²). This simplification allows little known but restricted species to fall in the “small” category even though their EOO was sometime originally extrapolated from a single location. Studies were geographically represented using ArcGIS (ESRI®). To compare mouse lemurs’ use of DF, we categorized the reported responses and/or use of DF into three categories: “positive effect” of forest degradation (mentions of preferential use of DF, higher abundance and greater fitness in DF), “neutral responses” (tolerance to DF, similar abundance at degraded and non-degraded sites, and foraging on cultivated plant species, or no detected differences), “negative responses” (exclusive use or higher abundance in primary forest, reduced fitness, reduced long-term population viability in DF, poaching, increased predation by domestic or wild animals in DF, and increased parasite spillover from humans or domestic animals). Single reports for observations of mouse lemurs in DF were not considered, since they do not indicate clear quantitative DF use trend.

To test for variation in degraded habitat use, with respect to forest type (dry, humid), conservation status and distribution range, we used a two sided Fisher’s exact independence test using R (R Core Team, 2015). This research adhered to the American Society of Primatologists’ principles for the ethical treatment of primates.

## RESULTS

We found a total of 84 studies (see the definition of “studies” in the method section) reporting effects of forest degradation on mouse lemurs. In 75 studies, the species names were specified (Tables 1, 2). In the other nine studies, the species name was not specified and could not be identified based on current taxonomy or geographic data (Table 2). Of these 75 studies, only 24 primarily focused on differential habitat quality use (*i.e.* 32%), but a larger proportion (n=65, *i.e.* 87%) evaluated responses of mouse lemurs towards DF (Table 2). Of these 65 studies, 27 (42%) reported negative, 23 (35%) neutral and 15 (23%) positive responses towards DF (Figure 1a, Tables 1, 2). While at the genus level a larger proportion of studies suggests that DF has a negative effect, our results also confirm that most of the studied mouse lemur species (12 out of 14) are able to use DF (Figure 1a, Table 2). However, reports of DF use are scarce for the majority of mouse lemur species, and unequally distributed across Madagascar. Most studies are concentrated in a few parks and sites with research facilities and long term research programs, such as Kirindy (9 studies), Ankarafantsika (8) and Ranomafana (13) (Figure 2), resulting in a paucity of data for numerous species outside of these parks. Hence, seven species are represented by one or two studies (Figure 1 main graph, Tables 1, 2) and ten species are not represented (e.g. *M. bongolavensis, M. jollyae*).

**Figure 1:**
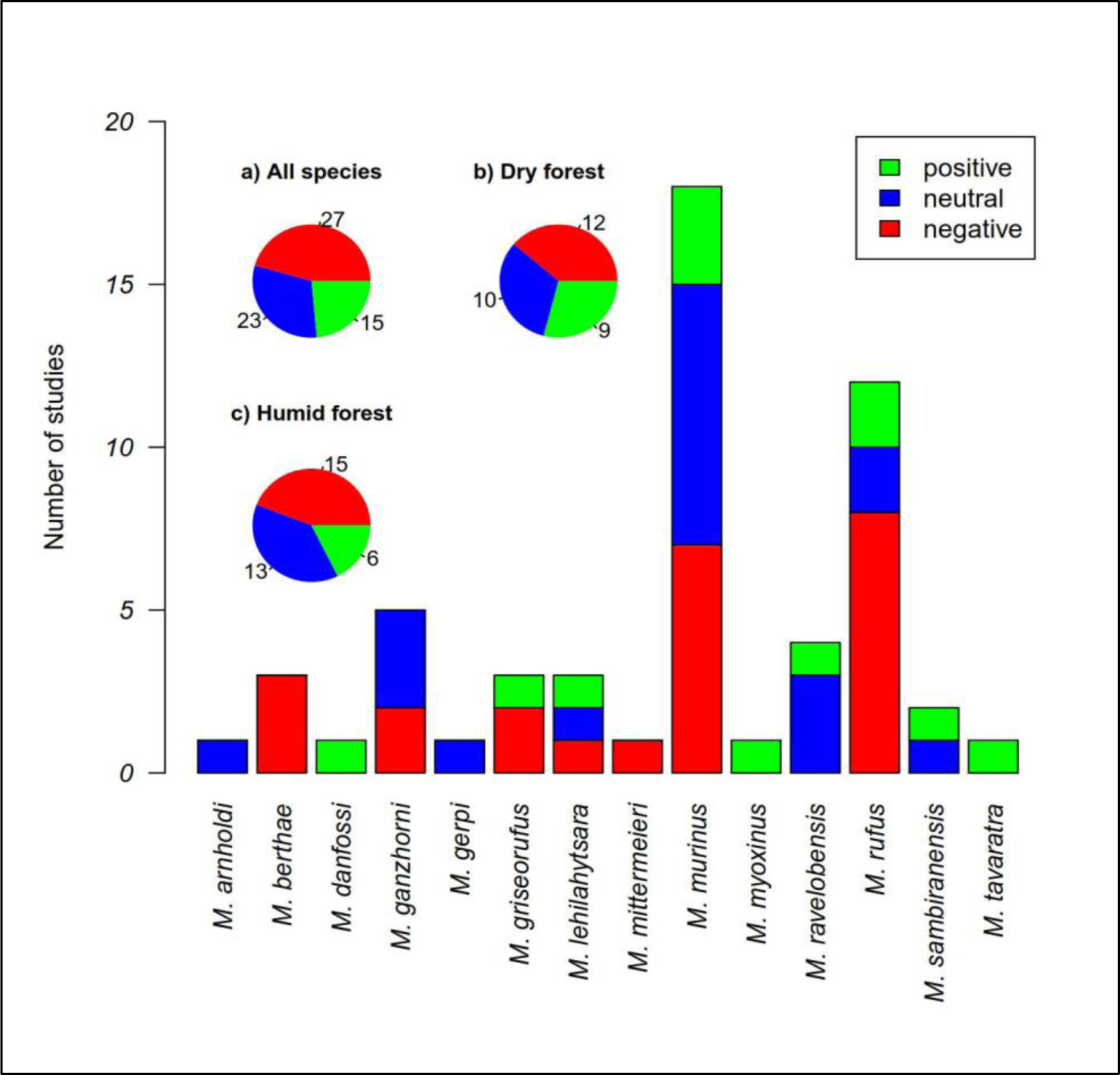
Forest degration effect on mouse lemurs. The main barplot represents the number of studies reporting negative, neutral or positive responses towards DF for each species. The pie charts represent the proportions (and numbers, beside the pie charts) of negative, neutral or positive responses obtained for **a)** all species, **b)** dry forest species and **c)** humid forest species.

**Figure 2:**
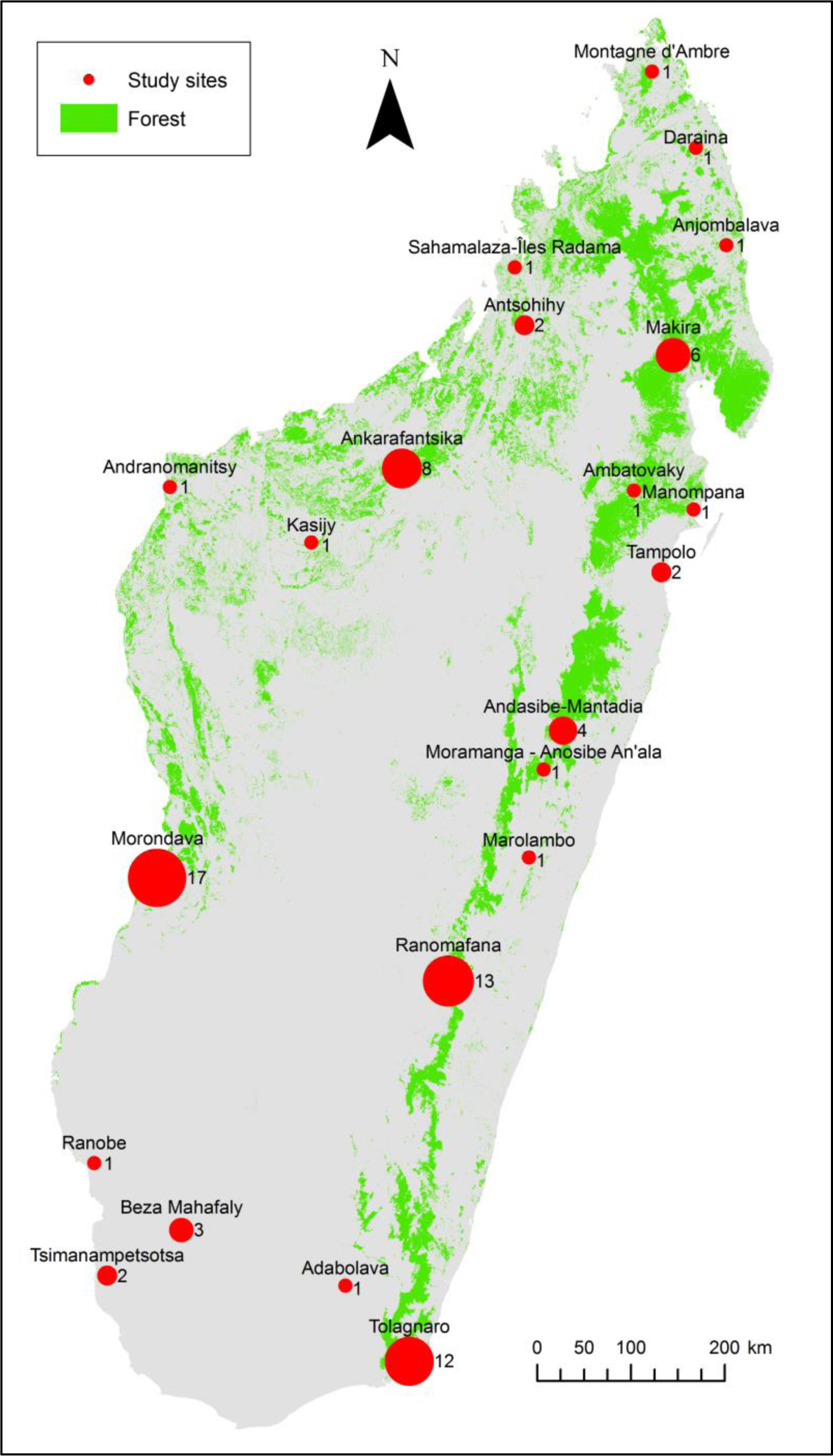
Geographic distribution of mouse lemur DF use studies. The diameter of the red dots are proportional to the number of studies (numbers beside dots) in the respective locations. Forest cover from the Madagascar Vegetation Mapping Project data (available online at <http://www.kew.org/gis/projects/mad_veg/datasets.html>; (Moat & Smith, 2007). Note that this figure represents numbers of single studies but the results description refers to “study” numbers as described in the method section.

**Table 1:** Summary of mouse lemurs’ degraded habitat use bibliography. Number of studies reporting use of degraded forest and factors invoked or demonstrated to influence degraded forest use per species. *NOTE*: ns= not specified. IUCN: conservation status: LC= Least Concern, VU= Vulnerable, EN= Endangered, CR= Critically Endangered. Dist. Range: L= large, M= medium, S= small. # Total: Number of studies reporting DF use, Response to DF= Number of studies assessing responses to DF. # Positive/ Neutral/ Negative= Number of studies reporting positive/ neutral/ negative responses. Positive/negative factors: main positive or negative factors invoke as influencing DF use. Comp= Competition. Diet= Diet (including insects and insect secretions, fruits, leaves, flowers and buds). Diet (F)= Diet (Fruits). D (I)= Diet (Insects). Predation (n)= Predation by native carnivores. Temp.= Temperature. a: Corbin and Schmid, 1995; Smith et al., 1997. b: Schäffler, 2011; Schäffler et al., 2015. c: Ganzhorn and Schmid, 1998. d: Raharivololona, 2009; Raharivololona and Ganzhorn, 2009. e: Gardner and Davies, 2014. f: Schmid, 1998. g: Andriamandimbiarisoa et al., 2015; Rakotondravony and Radespiel, 2009. h: Atsalis, 1999, i: Lehman, 2006; Lehman et al., 2006a; b; Rajaonson et al., 2010. j: Herrera et al., 2011. k: Wright et al., 2005; Karanewsky and Wright, 2015. l: Rasambainarivo et al., 2013; Bublitz et al., 2014; Zohdy et al., 2015. m: Ravoahangy et al., 2008; Lehman & Ratsimbazafy, 2001. n: Ratsirarson and Ranaivonasy, 2002; Goodman, 2003. o: Burke and Lehman, 2014. p: Rodriguez et al., 2015. q: Dammhahn and Kappeler, 2008a; b; 2009; 2010. r: Schwab and Ganzhorn, 2004. s: Ganzhorn, 1988. t: Ganzhorn, 1987. u: Ganzhorn, 1995. v: Meyler et al., 2012.

| Species | IUCN | Dist. Range | # Total | Response to DF | # Positive | # Neutral | # Negative | Positive factors | Negative factors |
| --- | --- | --- | --- | --- | --- | --- | --- | --- | --- |
| *M. murinus* | LC | L | 24 | 18 | 3 | 8 | 7 | Diet<sup>a</sup>, Predator release<sup>b</sup> | Diet<sup>c</sup>, Poaching<sup>be</sup>, Predation<sup>c</sup>, Tree holes<sup>cf</sup>, Temp.<sup>cf</sup>, Understory<sup>bc</sup> |
| *M. rufus* | VU | M | 14 | 12 | 2 | 2 | 8 | Diet<sup>hijk</sup>, Understory<sup>l</sup> | Diet<sup>k</sup>, Parasites<sup>l</sup>, Poaching<sup>lm</sup>, Predation<sup>n</sup>, Tree holes<sup>k</sup>, Temp.<sup>ik</sup> |
| *M. ganzhorni* | ns | ns | 7 | 5 | 0 | 3 | 2 | ns | Understoryg, Parasitesd |
| *M. ravelobensis* | EN | M | 7 | 4 | 1 | 3 | 0 | Diet (I)<sup>o</sup> | ns |
| *M. griseorufus* | LC | L | 8 | 3 | 1 | 0 | 2 | Parasites<sup>p</sup> | Poaching<sup>q</sup> |
| *M. berthae* | EN | S | 4 | 3 | 0 | 0 | 3 | na | Comp.<sup>brs</sup>, Understory<sup>bs</sup> |
| *M. lehilahytsara* | VU | M | 4 | 3 | 1 | 1 | 1 | Diet (F)<sup>t</sup> | Diet (F)<sup>u</sup>, Understory<sup>u</sup> |
| *M. sambiranensis* | EN | S | 2 | 2 | 1 | 1 | 0 | ns | ns |
| *M. arnholdi* | EN | S | 1 | 1 | 0 | 1 | 0 | ns | ns |
| *M. danfossi* | EN | M | 1 | 1 | 1 | 0 | 0 | ns | ns |
| *M. gerpi* | CR | S | 1 | 1 | 0 | 1 | 0 | ns | ns |
| *M. mittermeieri* | EN | S | 1 | 1 | 0 | 0 | 1 | ns | ns |
| *M. myoxinus* | VU | L | 1 | 1 | 1 | 0 | 0 | Diet (F)<sup>v</sup> | ns |
| *M. tavaratra* | VU | S | 1 | 1 | 1 | 0 | 0 | Understory<sup>w</sup> | ns |

**Table 2:**
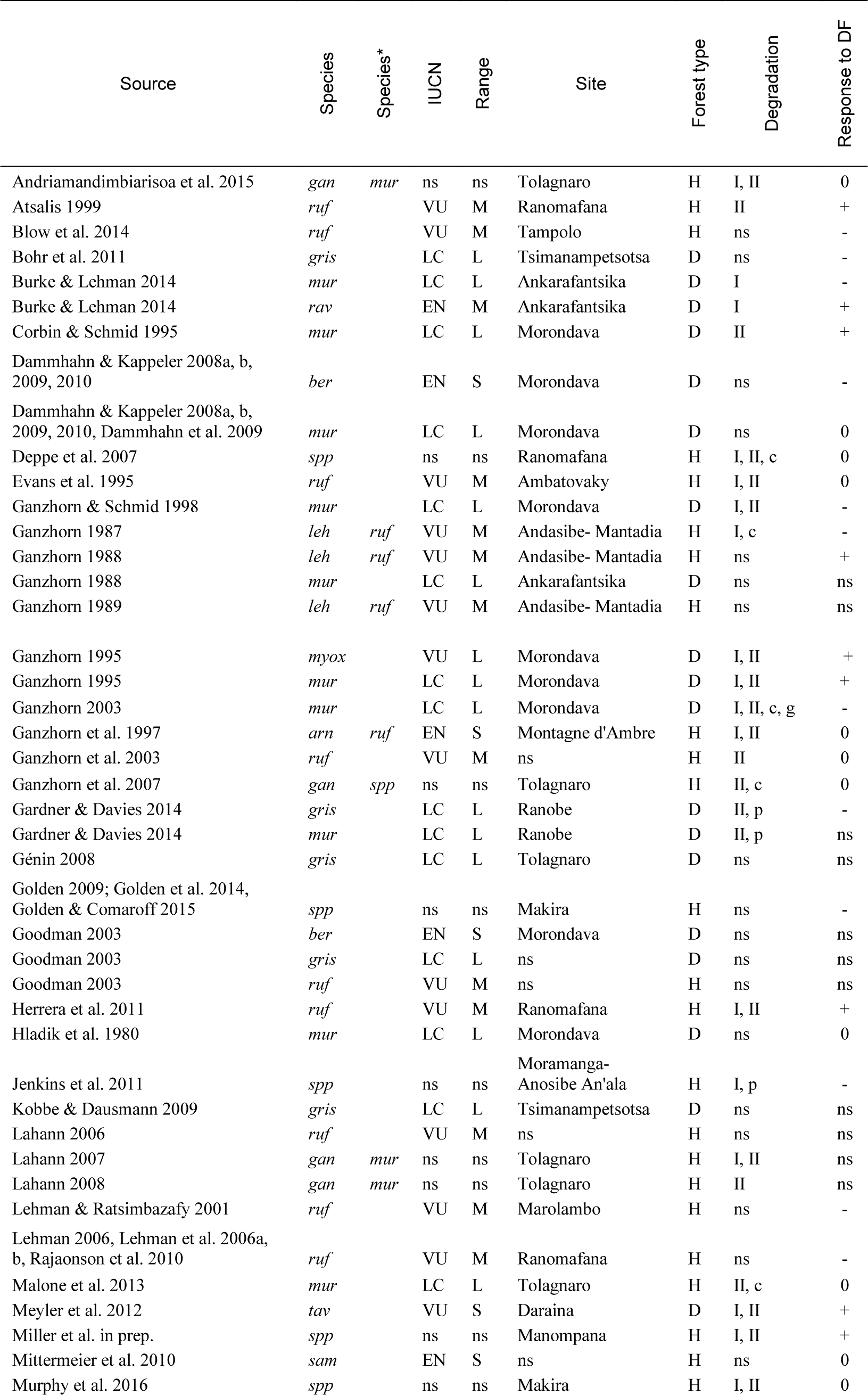
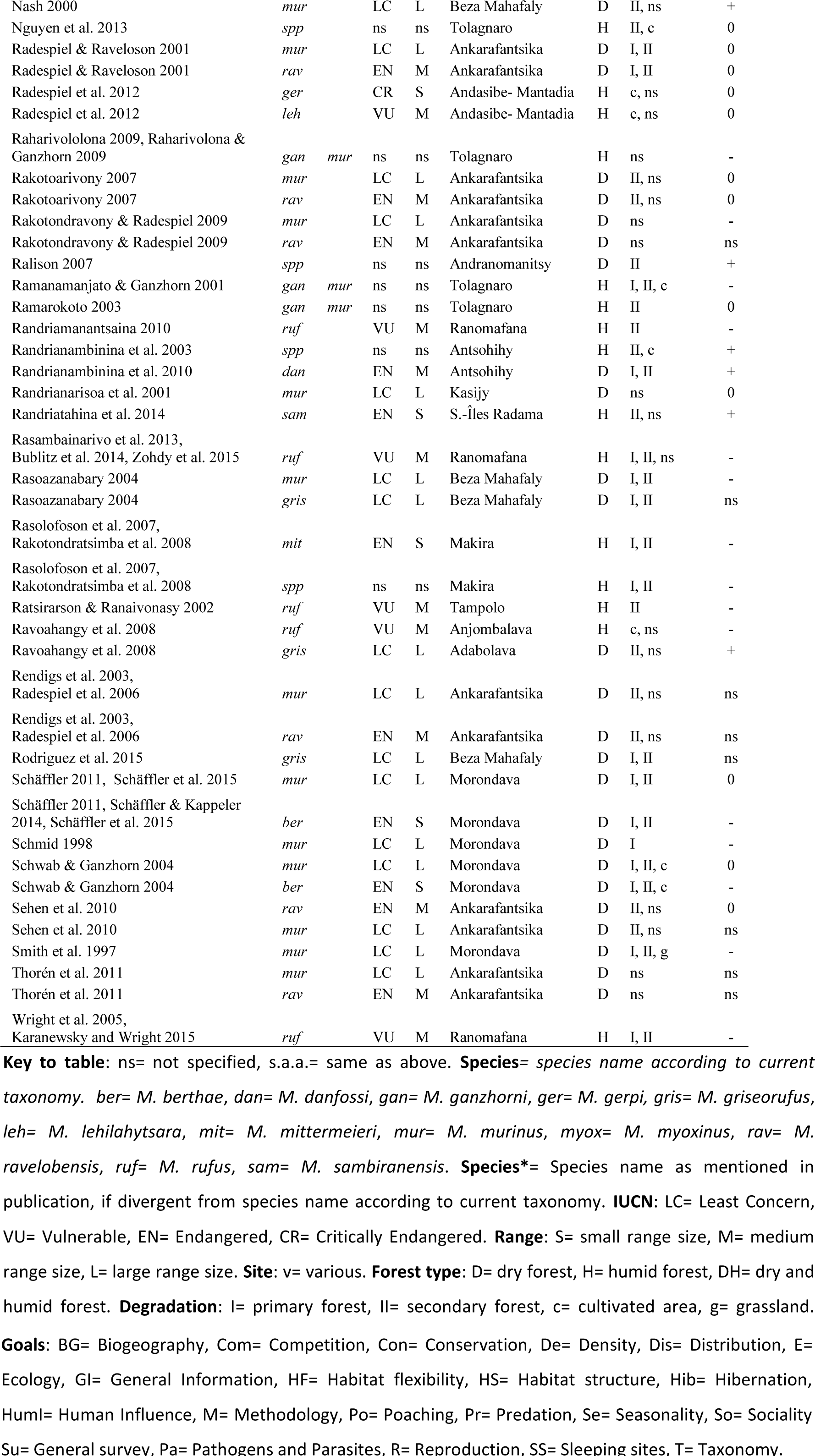
Overview on reviewed studies

Most species with at least three studies showed variable responses to DF with at least one positive effect report (Figure 1 main graph, Tables 1, 2). Similarly, the most frequently studied species, *M. murinus,* shows a high proportion of neutral responses (8 out of 18 studies) together with more negative than positive reports (7 vs. 3 studies). Likewise, *M. rufus* shows more negative than positive effects reports (8 vs. 2 studies). Of all species represented by more than two studies, *M. ravelobensis* (n=4) is the only one with no report of negative responses to DF (Figure 1 main graph, Table 1). In contrast, *M. berthae* was the only species for which only negative effects were reported (n=3, Figure 1 main graph, Table 1).

### Forest type

From the 65 studies evaluating responses towards DF, 31 were conducted in dry and 34 in humid forest (Figures 1b, c, Table 2). We found no difference in response to DF between dry and humid forests (Fisher’s exact test, *p*= 0.63). However, several reported or invoked factors potentially influencing response to DF showed contrasting frequency amongst forest types (Figure 3a). For instance, increased food availability was the most frequently mentioned reason for the use of DF (n=16) in both dry (n=8) and humid (n=8) forests, but the positive effects were associated to different causes. In dry forests, high insect abundance in degraded sites was invoked (n=6), whereas high fruit abundance in DF was invoked in humid forests (n=6); (Figure 3a).

**Figure 3:**
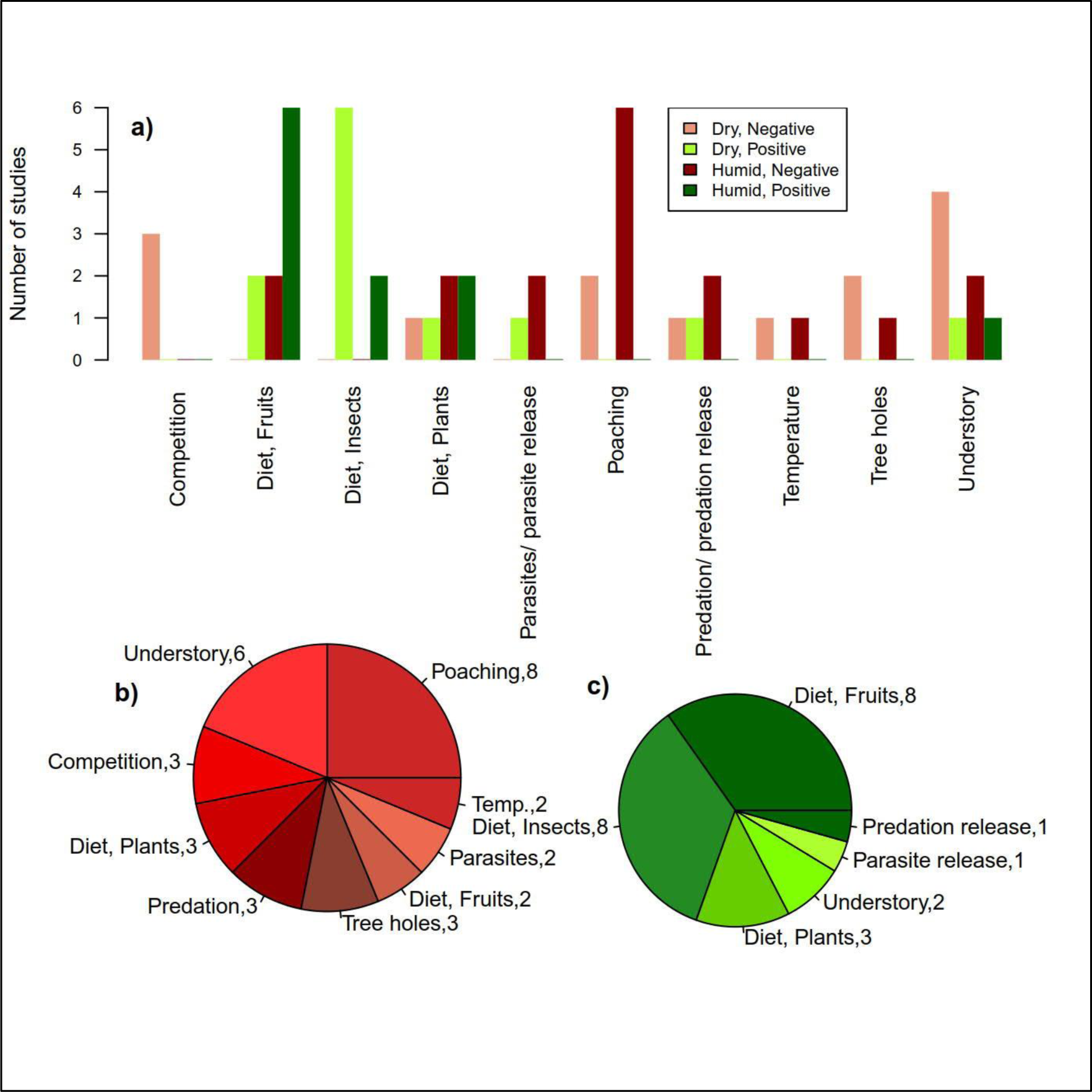
Reported or invoked causes of mouse lemur DF use. **a):** Numbers of studies suggesting positive or negative effects of factors on DF use in dry and humid forests, **b):** negative and **c):** positive effects reported to potentially influence mouse lemur use of DF. Pie sizes are proportional to the number of study.

### Distribution range size

Of the 14 species represented in the literature, six have a small, four a medium-sized, three a large, and one an undescribed distribution range (Tables 1, 2). We found a strong overrepresentation of species with large (n=22), and medium-sized (n=20) ranges and only a few studies (n=9) focusing on species with small distribution ranges (Figure 4a). In addition, we found no difference in responses to forest degradation amongst distribution range classes Fisher’s exact test, *p*=0.99, Figure 4a).

**Figure 4:**
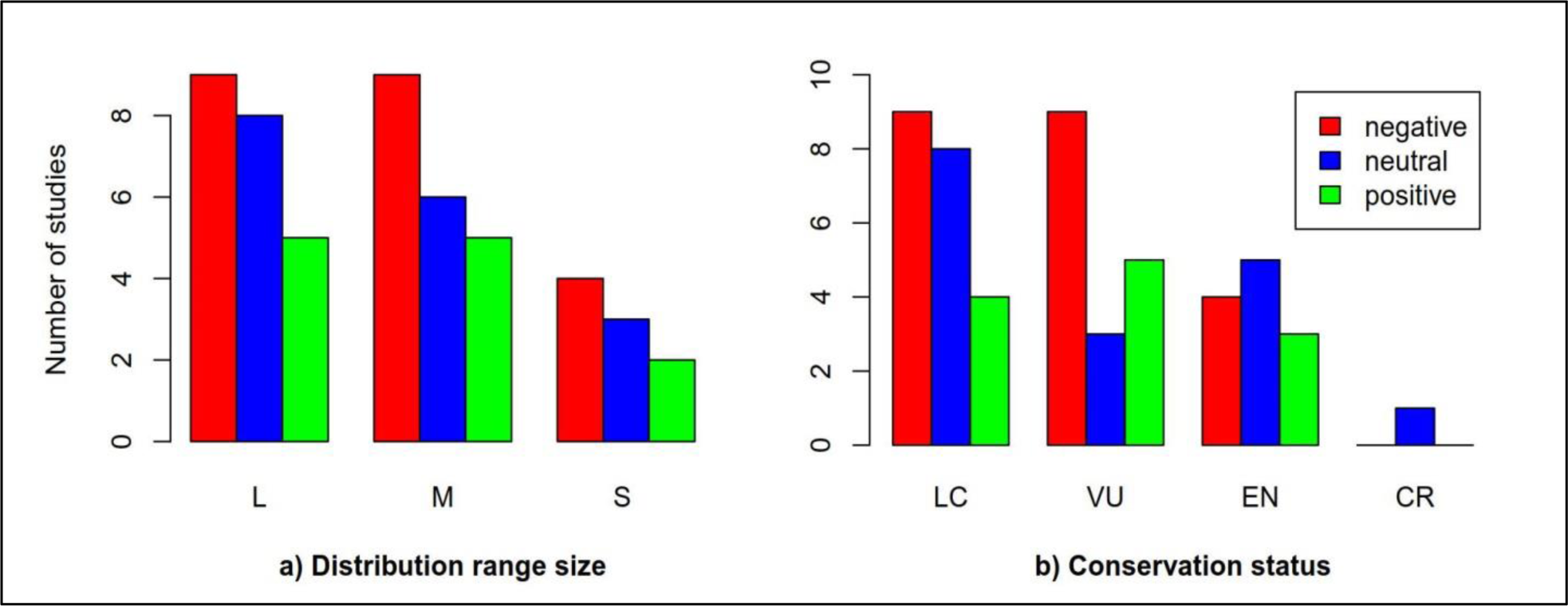
Relation between species distribution range size, conservation status and DF use. Number of studies reporting negative, neutral or positive effect of habitat degradation on mouse lemur use of DF are represented as function of their **a)** distribution range size (L: large, M: medium, S: small) and **b)** conservation status (LC: Least Concern, VU: Vulnerable, EN: Endangered, CR: Critically Endangered).

### Conservation status

The number of studies per species decreases with increasing conservation status (Figure 4b). Although 18 out of 24 lemur species are threatened (*i.e.* categorized as “Vulnerable”, “Endangered” or “Critically Endangered”) (IUCN, 2017; Schwitzer et al., 2013), there are almost as many studies on “Least Concern” species (n=32, most of them dealing with *M. murinus*) as on threatened species (n=37). Only one study focused on a Critically Endangered species, *M. gerpi* (Radespiel et al., 2012) (Figure 1 main graph, Figure 4b, Tables 1, 2). We found no significant difference in degraded habitat use between conservation status (Fisher’s exact test, *p*= 0.58), (Figure 4b). This may be due to the bias towards species with lower conservation status. Almeida-Rocha, Peres, & Oliveira (2017) found a similar pattern in a general pantropical meta-analysis of primates’ responses to DF.

### Factors potentially affecting the use of degraded forest habitats

Most of the 36 studies that invoke or investigate putative causes of DF use report food resources availability (44.4%; n= 16) and forest structure (30.6%; n= 11) as influencing mouse lemurs’ DF use. Poaching (22.2%; n= 8), predation (11.1%; n= 4), tree hole availability and pathogen transmission (8.3% each; n= 3) were also reported to potentially affect mouse lemur use of DF (Figures 3b, c, Table 1).

## DISCUSSION

Our literature analysis confirms that most mouse lemur species (12 out of 14) are able to use DF, even though they do not necessarily benefit from forest degradation. However, only 14 out of 24 species are represented in the literature and data on DF use is scarce for the majority of the represented species. In addition, it appears that most studies are concentrated in a few parks and sites with research facilities hosting long term research programs, as well as focused on a few overrepresented species (*M. murinus* and *M. rufus*). This unbalanced species representation and the overall low number of studies limited the power of our statistical inferences. However, it also highlights the lack of data for the most endangered (micro-endemic) species and stresses the need for a systematic and comprehensive investigation of species taxonomy, distribution, abundance and diet to accurately study mouse lemur use of DF (Lehman, Radespiel, & Zimmermann, 2016).

Despite dry and humid forest being substantially different and hosting mouse lemurs with distinct ecology (Kappeler & Rasoloarison, 2003; Radespiel, 2007) we found no clear variation of DF use between dry and humid forests (Q1). Nevertheless, the factors potentially influencing the use of DF varied (not statistically tested) between dry and humid forest. Food resources availability was reported or invoked by most studies investigating putative factors explaining the use of DF. However, while high insect abundance was positively associated with dry DF use, high fruit abundance was frequently reported from humid DF. This suggests that a systematic comprehensive investigation of diet in DF and non-DF is required to shed light on the differences between forests types as also suggested for the Cheirogaleidae family in general (Lehman et al., 2016).

Distribution range and conservation status (Q2 and Q3) are variables expected to be connected to habitat use flexibility. However, our analyses of the literature showed no evidence of relation between distribution range, conservation status and DF use. However, it should be kept in mind that most mouse lemur species have been described in the last decades (Hotaling et al., 2016; Radespiel et al., 2012; Rasoloarison et al., 2013) and both their taxonomy and distribution range are not yet fully and definitively characterized (Hotaling et al., 2016; IUCN, 2017; Lehman et al., 2016; Louis Jr. & Lei, 2016; Schwitzer et al., 2013). Therefore, relationship patterns between these variables and the use of DF might emerge in the near future from the completion of these data-sets. The conservation status is a complex and frequently evolving variable influenced not only by the distribution range and its variation but also by the species demographic trends and by the development of threats (IUCN, 2012). Hence, it is not necessarily surprising that we could not find a clear relation between DF use and the conservation status. In addition, our review highlights that food resources availability and habitat structure (e.g. understory) are the main factors invoked and/or reported to influence DF use. Below, we further discuss major putative factors in greater detail and finally propose a systematic and comprehensive framework to investigate DF use patterns.

### Food resources availability

Food resources availability was the most frequently invoked factor to explain differential use of DF (Figure 3) and is seen by many authors as a decisive factor determining the survival (Ganzhorn & Schmid, 1998; Hladik, Charles-Dominique, & Petter, 1980), the abundance (Bohr, Giertz, Ratovonamana, & Ganzhorn, 2011; Ganzhorn, 1988; Lehman, Rajaonson, & Day, 2006a; Sehen et al., 2010), and the reproductive success (Wright, Razafindratsita, Pochron, & Jernvall, 2005) of mouse lemurs. Although mouse lemurs are omnivorous (Mittermeier et al., 2010), their diet varies amongst species and seasons (Dammhahn & Kappeler, 2008, 2009; Radespiel, Reimann, Rahelinirina, & Zimmermann, 2006; Rakotondranary, Struck, Knoblauch, & Ganzhorn, 2011; Thorén et al., 2011). A large number of studies (n=16) invoked or reported higher abundance of particular food resources in degraded forests (Figure 3, Table 1). One of the most frequently invoked or reported positive effect of forest degradation is the abundance of insects in DF and along forest edges (Figure 3, Table 1), which constitute a considerable share of several mouse lemurs species’ diet (Corbin & Schmid, 1995; Lehman et al., 2006a). Finally, mouse lemurs have been reported to feed on cultivated plant species (Deppe et al., 2007; Ganzhorn, Goodman, & Dehgan, 2003), further emphasizing the role of mouse lemur diet flexibility for its use of modified habitat. However, negative effects were suggested, often by the same authors. For instance, Wright et al. (2005) pointed out that a large number of tree species selectively logged for wood are important components of *M. rufus’* diet.

### Understory structure and tree hole availability

Mouse lemurs are mostly found in the shrub and understory layer of the forest (Hladik et al., 1980; Kappeler & Rasoloarison, 2003). A dense understory seems to constitute the ideal substrate for feeding (Andriamandimbiarisoa et al., 2015; Radespiel et al., 2006), sleeping (Rasoazanabary, 2004), movements and locomotion (Andriamandimbiarisoa et al., 2015; Ganzhorn, 1987). Although anthropogenic disturbances may have a negative effect on understory structure, several authors reported positive selective logging and degradation effects on understory plant production and density (Ganzhorn, 1995, 1999; Herrera et al., 2011). For instance, Miller et al. (forthcoming) found higher population densities in the dense understory of mature secondary forest. Similarly, Ganzhorn (1987) reported the presence of mouse lemurs in old (but not young) *Eucalyptus* plantations with a developed shrub layer.

Tree holes constitute ideal shelters for daily torpor, sleeping, communal breeding and against predation for hollow dwelling species (Ganzhorn & Schmid, 1998; Karanewsky & Wright, 2015; Radespiel, Zimmermann, & Jurić, 2009). Selectively logged or degraded forests may provide less suitable tree hole shelters (Figure 3), a potentially limiting resource for hollow dwelling mouse lemurs’ DF use, in times of resource scarcity and climatic extremes (Ganzhorn & Schmid, 1998; Karanewsky & Wright, 2015; Kobbe & Dausmann, 2009; Schmid, 1998).

### Predation and poaching

Poaching pressure is often associated with DF and forest edges (Lehman, Rajaonson, & Day, 2006b; Lehman & Wright, 2000). Eight studies negatively associated mouse lemur poaching with differential use of DF (Figure 3, Table 1). Although mouse lemurs suffer lower hunting pressure than larger-bodied lemur species (Jenkins et al., 2011; Lehman & Ratsimbazafy, 2001), they are consumed by humans (Gardner & Davies, 2014; Jenkins et al., 2011). In addition, domestic carnivores (*Canis familiaris)* (Gerber, Karpanty, & Randrianantenaina, 2012; Goodman, 2003) and *Felis catus* (Gerber et al., 2012; Ratsirarson & Ranaivonasy, 2002) are likely to forage more frequently along forest edges (Figure 3, Table 1) and in forests used by humans (Farris, Gerber, et al., 2015; Farris, Golden, et al., 2015). Contrastingly, mouse lemurs may reduce predation rates from wild predators (carnivores, snakes) (Goodman, 2003; Ratsirarson & Ranaivonasy, 2002), birds of prey (Goodman, 2003; Mittermeier et al., 2010) by foraging in dense understory vegetation and by resting in tree holes (Rasoazanabary, 2004; Schmid, 1998). Indeed, higher predation pressure in DF was used to explain low DF use in three studies (Figure 3, Table 1). Contrastingly, Schäffler et al. (2015) suggested a positive effect of predation on DF use (decreased predation of *M. murinus* by *Mirza spp*.), which in turn released *M. berthae* from competition in primary forest.

### Conservation Implications

We highlight five factors frequently reported or invoked as influencing DF use: (i) food resources availability, (ii) understory and forest structure, (iii) poaching and predation, (iv) tree hole availability and (v) pathogen transmission. Besides the work required to limit or stop deforestation, forest degradation and poaching, namely the most important threats to lemur populations (IUCN, 2017; Schwitzer et al., 2013; Schwitzer, Mittermeier, et al., 2014), conservation managers may need to consider these five factors (also highlighted in Lehman et al. (2016)). For instance, reforestation projects may want to consider plant species belonging to the diet of mouse lemurs (and other species) such as *Bakerella* spp. (Atsalis, 1999), fruit trees (Atsalis, 1999; Ganzhorn, 1988), trees favoring high insect abundance, as well as hollow-forming trees (e.g. *Strychnos madagascariensis* (Salmona et al., 2015), and fast growing shrubs to facilitate dispersal and provide shelter for mouse lemurs (Andriamandimbiarisoa et al., 2015). Furthermore, conservation projects considering practices beneficial to rural communities and wild populations may carefully weigh the effect of selective logging and poaching. Conservation projects including localized selective logging (e.g. “KoloAla Manompana” (Rakotomavo, 2009)) may not be detrimental to mouse lemur populations (Atsalis, 1999; Ganzhorn, 1995), if middle sized trees, the understory and the shrub layer are maintained. In addition, although several studies reported mouse lemurs’ poaching (Gardner & Davies, 2014; Jenkins et al., 2011) and its negative effects on DF use (Figure 3, Table 1), it seems not be as frequent as for larger-bodied lemur species (Jenkins et al., 2011; Lehman & Ratsimbazafy, 2001). Mouse lemur populations are likely to be less susceptible to poaching than larger-bodied lemurs because of their shorter generation time and higher reproductive rate (Hohenbrink, Zimmermann, & Radespiel, 2015; Zimmermann & Radespiel, 2013). Therefore, mouse lemur harvesting needs to be formally evaluated to determine under which conditions sustainability can be achieved (Gardner & Davies, 2014; Golden, 2009).

## CONCLUSION

Our literature review analysis highlights that although most mouse lemur species are able to use DF, they are not necessarily favored by DF. Furthermore, it sheds light on the fact that data on DF use is geographically aggregated in a few locations (Figure 2), lacking for half of the described species and scarce for the majority of others. This stresses the need for a systematic and comprehensive investigation that will allow to accurately quantify the use of DF across species and regions. Field efforts should aim at comparing multiple species, and focus on filling the existing data gap for most micro-endemic species. They should combine density estimates methods such as nocturnal distance sampling and capture mark recapture (e.g. (Meyler et al., 2012)), with habitat characterization and opportunistic fecal material collection. In particular, habitat characterization may focus on describing forest structure (Lehman, 2016), flora and fauna diversity, but also on predator abundance using camera traps (e.g. (Farris, Gerber, et al., 2015; Farris, Golden, et al., 2015)) and tree hole availability. In addition, opportunistic fecal material sampling from capture studies combined with emergent meta-barcoding approaches will bring a better understanding of diet and parasite load (De Barba et al., 2014; Quéméré et al., 2013) in complement to arduous field observations. Finally, combined continuous field and genetic efforts (Hotaling et al., 2016; Louis Jr. & Lei, 2016; Yoder et al., 2016) will likely bring soon an accurate representation of species distribution and taxonomy necessary to study such ecological patterns at the genus scale. While our work focused on mouse lemurs, the second most speciose lemur genus, we stress that DF use should be studied across vertebrate species. In fact, similar studies will be required across all animals, plants and fungi as most habitats are likely to become increasingly fragmented and degraded in the future.

## ACKNOWLEDGEMENTS AND FUNDING INFORMATION

We thank Ute Radespiel, Isa A. Pais and Gabriele Sgarlata for comments and discussion on early versions of the manuscript as well as a large number of anonymous reviewers that helped substantially improving the manuscript. Financial support for this study was provided by the ‘Fundação para a Ciência e a Tecnologia’ (grant number SFRH/BD/64875/2009 to J.S. and grant numbers Biodiversa/0003/2015, PTDC/BIA-BIC/4476/2012, PTDC/BIA-BEC/100176/2008 to L.C.), the GDRI Madagascar, the ‘Laboratoire d’Excellence’ (LABEX) entitled TULIP (ANR-10-LABX-41), ‘Rufford Small Grant Foundation’ (grant number 10941-1 to J.S.), the Instituto Gulbenkian de Ciência, the LIA BEEG-B (Laboratoire International Associé - Bioinformatics, Ecology, Evolution, Genomics and Behaviour) (CNRS). This study was conducted in agreement with the laws of the countries of Portugal, France and Madagascar.

